# Species-specific phyllosphere responses to external pH change

**DOI:** 10.1101/2024.11.01.621601

**Authors:** Cristal Lopez-Gonzalez, Jean-Baptiste Floc’h, Tanya Renner, Kadeem J. Gilbert

## Abstract

The leaf surface, known as the phylloplane, represents the initial point of contact for plants in their interaction with the aboveground environment. Although prior research has assessed how leaves respond to external pH variations, particularly in the context of acid rain, there remains a limited understanding of the molecular mechanisms through which plants detect, respond to, and mitigate cellular damage. To look at plant responses to external pH changes, we measured the phylloplane pH for five species with variable phylloplane pH that ranged in the dry control. Moreover, we investigated the phylloplane pH in response to three pH treatments (pH 6.5, 4, and 2) and found that plants can modify their phylloplane pH, and this buffering ability is species-specific. Among the species analyzed, only *Gossypium* displayed a strong buffering ability. For treatments where leaves were exposed to either pH 6.5 or pH 4, *Gossypium* alkalinized the phylloplane pH slightly higher than the dry control pH. Remarkably, when leaves were exposed to pH 2, *Gossypium* was able to buffer the pH to 6 within five minutes. Furthermore, our transcriptional analysis indicated that the responses to external pH changes varied among species, highlighting differentially expressed genes associated with calcium (Ca^2+^) signaling pathways, as well as Ca^2+^ and H^+^-ATPases pumps. These findings also suggest that pH stress negatively impacts photosynthesis, and that both wetness and moderate pH shifts may trigger additional abiotic and biotic stress signaling pathways.

## Introduction

The phylloplane, i.e., the outermost surface of a leaf, is the initial point of interface between plant aboveground tissues and the external environment (Gilbert and Renner, 2021; Shepherd and Wagner, 2007). Plants must face a multitude of external abiotic environmental parameters on these surfaces, such as temperature and moisture (Kashyap et al., 2021; Munns and Millar, 2023; Pereira, 2016; Zhang et al., 2022, 2023). How physiological changes in evapotranspiration alter leaf temperature has been investigated, including the ecological consequences of this leaf microenvironment heterogeneity (Griffani et al., 2024; Kibler et al., 2023; Xue et al., 2020). Another parameter of likely importance is pH. The pH influences the chemical reactions that occur in multiple systems (Tsai and Schmidt, 2021a). Leaves may regularly experience external changes in pH, either through precipitation or other fluids that come into contact with the leaf in nature, and it would therefore be advantageous for plants to be able to sense external pH changes and appropriately respond to counter harmful changes (Cornelissen et al., 2006; Cornelissen et al., 2011; Tao et al., 2019; Tsai and Schmidt, 2021; Rodríguez-Sánchez et al., 2020). The ability of plants to control pH is studied mainly in the rhizosphere (Husson, 2013; Schmidt, 2022). In the rhizosphere, soil pH can control the availability of mineral nutrient absorption, the structure of the soil microbiome, and other plant abiotic and biotic interactions (Husson, 2013; Lammel et al., 2018; Muneer et al., 2021; Schmidt, 2022; Tsai and Schmidt, 2021b). Given its relevance to the rhizosphere, the regulation of external pH should also be important to study in the phylloplane. Based on the current literature, the phylloplane pH can play a role in how plants mitigate acid rain, fertilizers and pesticides sprayed onto the leaf, and potential plant-microbe interactions (Lager et al., 2010; Perreault and Laforest-Lapointe, 2022; Shepherd and Wagner, 2007; Shi et al., 2021; Simon et al., 1994; Tao et al., 2019; Wong, 1997). Furthermore, the phylloplane pH can vary between lineages and species and is influenced by moisture (Cornelissen et al., 2011; Gilbert and Renner, 2021). Most plants tend to have a neutral pH, such as *Beta vulgaris*. Only species belonging to the Malvaceae family have been reported to have a highly alkaline phylloplane pH under normal conditions (Gilbert and Renner, 2021; Harr et al., 1980); these pH levels increase further after spraying water on their leaf surface (Harr et al., 1980; Navon et al., 1988; Smith et al., 1996). The opposite extreme of the phylloplane pH is observed in carnivorous plants, ranging from below pH 5 to extremely low pH 1 (Gilbert, 2020; Gilbert and Renner, 2021).

Simulated acid rain is the most common type of experiment related to phylloplane pH disturbance. These studies have shown that leaves can neutralize acidic pH and mitigate foliar damage (Adams and Hutchinson, 1984; Knittel and Pell, 1991; Liang et al., 2015; Ren et al., 2018; Rodríguez-Sánchez et al., 2020; Shi et al., 2021).

Additionally, this buffering ability is species-specific, and its effectiveness depends on the acidity and duration of the external disturbance (Adams & Hutchinson, 1984; Liang et al., 2015; Liu et al., 2013; Pfanz & Heber, 1986; Ren et al., 2018; Zheng et al., 2017). Consequently, this ability to actively regulate phylloplane pH might have consequences for how the leaf surface interacts with biotic and abiotic stressors, such as microbial interactions (Perreault and Laforest-Lapointe, 2022; Schmidt, 2022), and thus, can play a role in the fitness and productivity of agricultural crops (Bashir et al., 2022). Despite this, the molecular mechanisms behind the plant response to pH as a stressor have not been studied in detail. Fundamental questions remain unanswered, including whether there are any specific pathways involved in this response or if this mechanism is converging with other abiotic stress, or whether the primary mechanism is shared across plant taxa.

Here, we use a comparative transcriptomic approach to elucidate the key genes involved in pH buffering ability. We have selected an assemblage of plants with known variation in pH ranging from alkaline to acidic extremes (*Gossypium arboreum*, *Gossypium hirsutum*, *Beta vulgaris*, *Nepenthes bicalcarata,* and *Nepenthes rafflesiana*). We tested a 5-minute response to acidic pH disturbance in the leaf blade due to the rapid ability of plants to buffer pH. Only a few studies have reported fast transcriptional reprogramming responses to an external stimulus such as wounding and heat stress (Hander et al., 2019; Lohani et al., 2022). In addition, studies documenting these rapid responses to stress are needed to understand how plants manage to adapt to a changing environment. Here, we found that plant response to pH disturbance is interlinked to the general signaling pathway involved in other abiotic stresses and that plant response to pH disturbance is species-specific.

## Results

### Analysis of pH regulation ability

We selected five different species with predicted pH levels ranging from alkaline to acidic to assess a broad initial phylloplane pH: *Gossypium arboreum*, *Gossypium hirsutum*, *Beta vulgaris*, *Nepenthes bicalcarata,* and *Nepenthes rafflesiana*. For each species, we exposed experimental leaves to one of three pH treatments (6.5, 4, 2) for 5 minutes and some were left unsprayed as dry controls. We then measured their phylloplane pH levels to test the buffering ability of plants in response to external pH changes.

Both *Gossypium* species had an alkaline initial (dry, unsprayed) phylloplane pH (*G. arboreum*, 8.7 ± 0.1; *G. hirsutum*, 9.0 ± 0.4), while *Beta vulgaris* was nearly neutral (7.4 ± 0.1) and both *Nepenthes species* were more acidic (*N. bicalcarata*, 4.8 ± 0.3; *N. rafflesiana*, 5.5 ± 0.6; Figure 1A). We also examined the resultant pH levels the leaves achieved in response to each treatment. The resultant pH was significantly different between each genus compared to *Beta vulgaris* (Supp. Table 1). Interestingly, within each species, the resultant pH levels were similar for the Wet (i.e., neutral control) and pH 4 treatments (Figure 1A). This shows that each species was able to effectively buffer their pH under pH 4 treatment (i.e., return to normal baseline levels). *N. bicalcarata* acidified its phylloplane in response to both Wet and pH 4, achieving pH 3.96 ± 0.6 and 3.62 ± 0.4, respectively (Figure 1A, Supp. Table 1). Although the resultant pH levels did not differ between Wet (p-value = 0.59) and pH 4 (p-value = 0.65) conditions within species (Supplementary Table 1), they did vary across species/genera (p-value < 0.001, Figure 1A, Supplementary Table 1).

**Figure 1.**
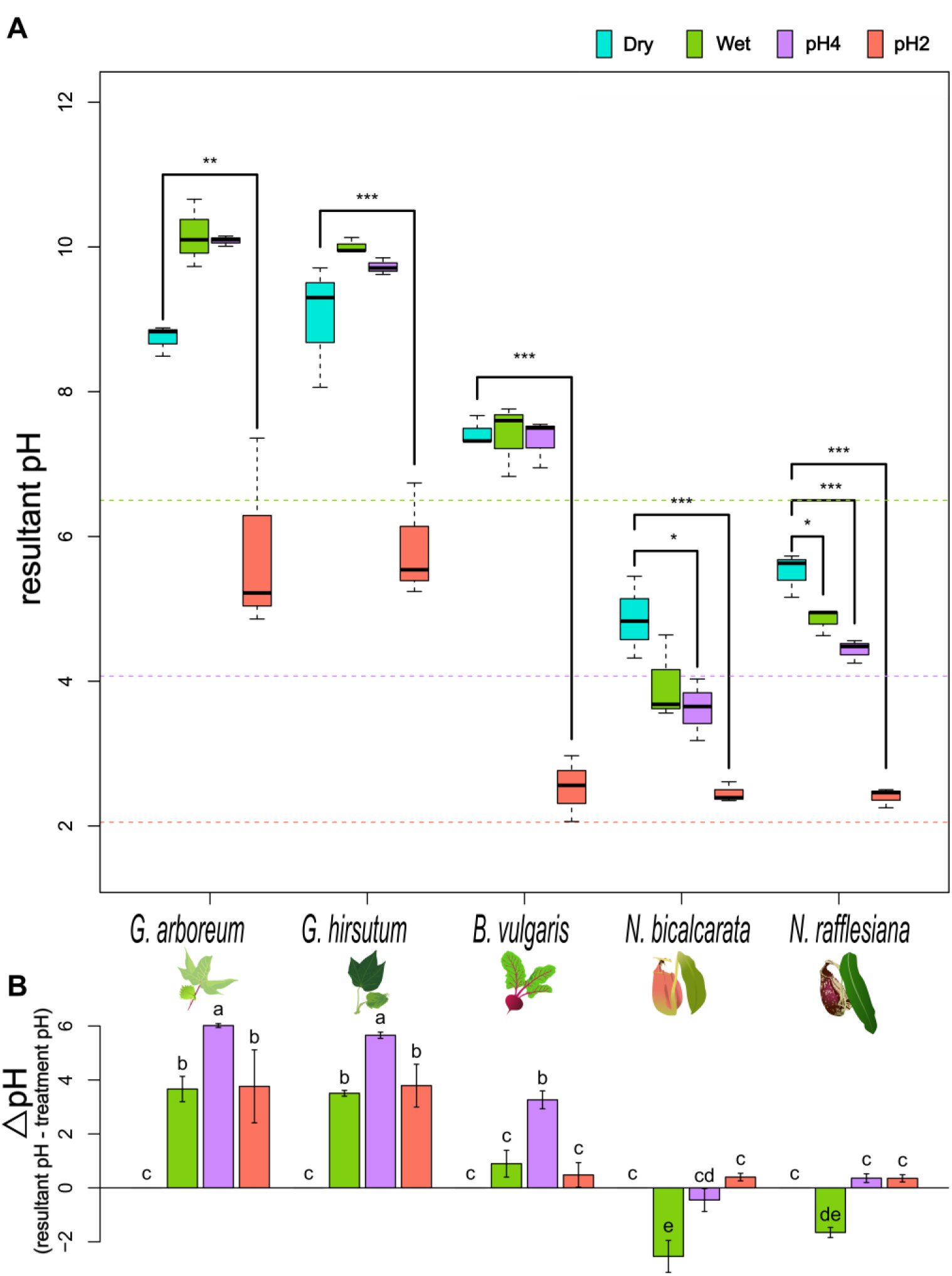
Buffering responses to external pH changes differ by species. A) The dry phylloplane pH across different species and 5 min after applying one of the three pH treatments, Wet (pH 6.5; green dotted line), pH 4 (purple dotted line) or pH 2 (orange dotted line) treatment. Brackets indicate significant differences between treatments; Dunnet test, n=3, p-value indicated by number of asterisks; * 0.05, ** 0.01, *** 0.001. B) Buffering response, i.e., the calculated difference between the resultant pH and treatment pH for each species; letters indicate differences across species LSD-test, n=3. Delta pH for Wet treatment indicates *Gossypium* species are alkalinizing (positive values), *Beta* is neutral (not different from zero) and *Nepenthes* are acidifying (negative values). Delta pH for the pH4 treatment shows both *Gossypium* and *Beta* are buffering (different from zero) whereas *Nepenthes* are not. Delta pH for pH2 treatment shows that only *Gossypium* species buffered against this treatment.

Certain pH response patterns at the genus level were observed. The *Gossypium* species increased pH slightly higher than the dry control under both Wet and pH4 treatments (pH 9.6-10.7); even when not significant with respect to dry pH, it exhibited a similar pattern in both *Gossypium* species. *Beta vulgaris* buffered both Wet and pH4 to similar values relative to the dry control. Finally, both *Nepenthes* species had different responses from *Gossypium*: they slightly acidified their pH under Wet and pH4 treatments; up to a ∼2 unit pH difference in the former case. In all genera, pH2 treatment significantly impacted the resultant pH (Supplementary Table 1). However, only *Gossypium* was able to significantly buffer against pH2 treatment, showing on average a ∼4 unit increase in pH at the end of the 5-minute response period (Figure 1B). Buffering responses varied across species and were pH-dependent.

### Differential gene expression analysis

To identify key genes involved in the response to external pH changes, we performed an RNAseq analysis using our experimental leaves. After filtering transcripts by low counts (< 3 TPM) and selecting only the most expressed transcript per gene, we ended up with 96,097 transcripts (13.064%) for *G. arboreum*, 64,429 transcripts (16.93%) for *G. hirsutum*, 53,788 transcripts (12.35%) for *B. vulgaris*, 85,027 transcripts for *N. bicalcarata* and 124,547 transcripts for *N. rafflesiana*. We annotated genes using EnTAP (Hart et al., 2020) and selected only genes annotated as Viridiplantae. The number of genes remaining were 16, 742 in *G. arboreum*, 17, 239 in *G. hirsutum*, 11, 274 in *B. vulgaris*, 12, 719 in *N. bicalcarata,* and 13, 325 in *N. rafflesiana*.

We conducted differential gene expression (DGE) analysis with Deseq2 (Emms and Kelly, 2019; Love et al., 2017) using a multifactorial array consisting of 1) comparing each pH treatment (Wet, pH4, pH2) against the dry control (WetvsDry, pH4vsDry and pH2vsDry) in order to elucidate genes involved in the general disturbance to wetness and pH; and, 2) comparing pH4 and pH2 against the Wet (pH 6.5) treatment (pH4vsWet and pH2vsWet) to identify genes that could be more directly related to pH changes (Figure 2). We identified a total of 279 DEGs for *G. arboreum*, 290 DEGs for *G. hirsutum,* 90 DEGs for *B. vulgaris,* 567 DEGs for *N. bicalcarata,* 358 DEGs for *N. rafflesiana* in all treatment comparisons, with a cut-off parameter of log-fold change (logFC) ≥ |1| and false discovery rate (FDR) < 0.05. We examined the number of DEGs for each comparison and each species (Figure 2). Figure 2 Each species showed a different number of DEGs across treatment comparisons (Figure 2); *B. vulgaris* was the species with the fewest DEGs in general and across treatment comparisons; pH6vsDry (55 DEGs) was the one comparison that contained more than half the total DEGs found in this species. Interestingly, *N. bicalcarata* showed the highest number of DEGs in all the species, and the pH4vsDry comparison (368 DEGs) seems to drive more than half of them.

**Figure 2.**
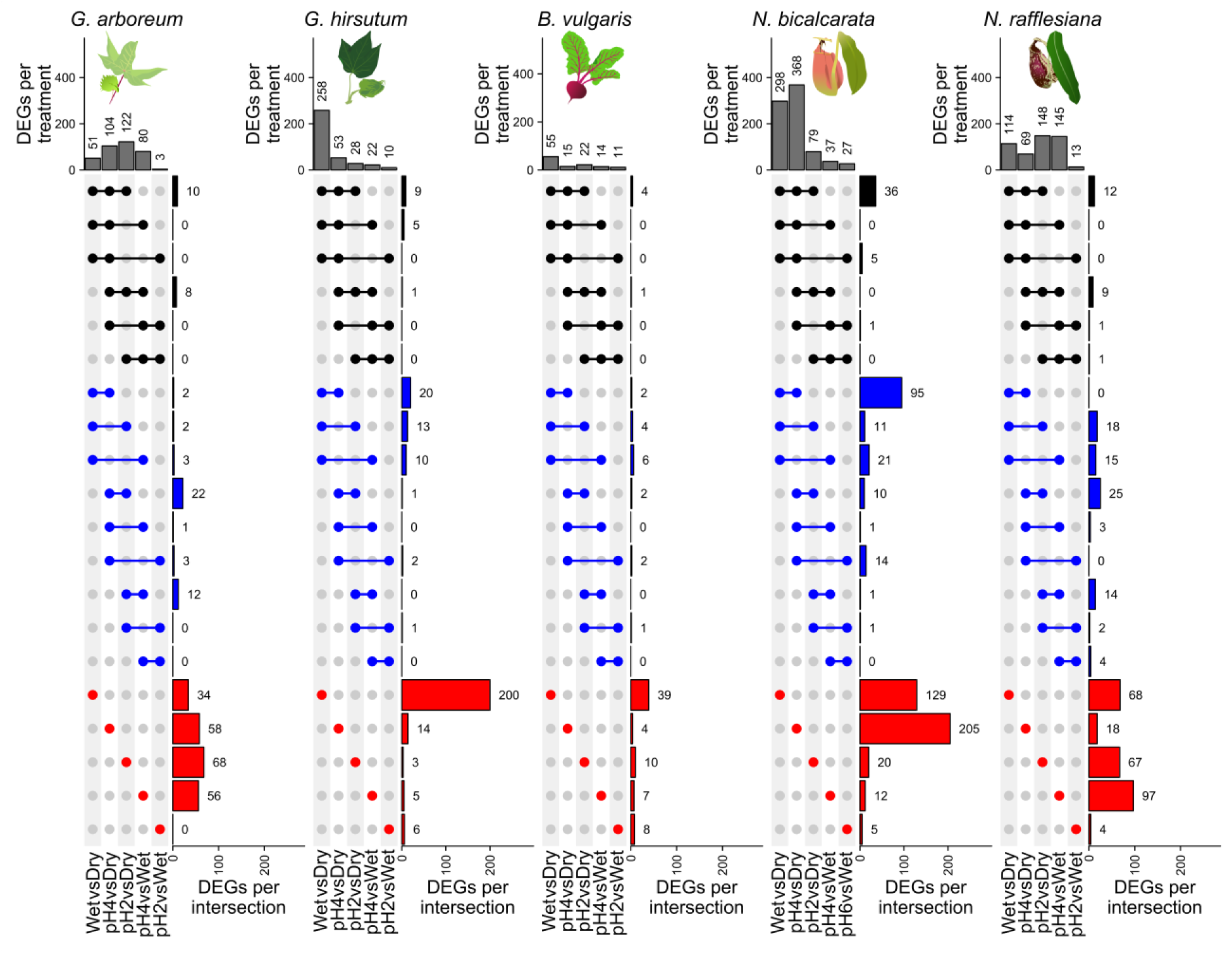
The number of differentially expressed genes (DEGs) in response to external pH changes varies by species and treatment comparison. The number of DEGs are summarized in the top bar plot per species. Colors indicate the number of intersections across treatments: red for unique singletons, blue for pairwise comparisons, and black for three-way comparisons. DEGs were chosen with a cutoff of log Fold Change > |1| and p-adjusted value < 0.05. Beta vulgaris appears to be the least responsive species to external pH change by the number of DEGs. In contrast, N. bicalcarata has 567 total DEGs and has 205 DEGs uniquely expressed in pH4vsDry treatment therefore this species appears to be more sensitive to environmental changes.

Most DEGs are uniquely expressed in only one treatment comparison, with fewer genes shared between two- or three-treatment comparisons (Figure 2). Genes involved in a general response to external disturbance and wetness were expected in the genes shared in all three comparisons of wet, pH4, and pH2 against dry control (Figure 2, black arrow); while genes related to pH responses were expected in the genes shared in the pH4vsDry and pH2vsDry comparison, and pH4vsWet and pH2vsWet comparison (Figure 2) and uniquely differentially expressed in the pH2vsWet comparison; but no clear pattern was observed (Supplementary Data). Both species of *Gossypium* had a similar number of total DEGs, but the distribution of DEGs in treatment comparisons was quite different (Figure 2). The most drastic difference was observed in *G. hirsutum*, where pH6vsDry comparison (258 DEGs) showed the most DEGs and they were mostly uniquely expressed (200 DEGs) in that treatment comparison (Figure 2).

### Orthologues analysis

We ran OrthoFinder (Emms and Kelly, 2019) to determine which genes are orthologues across species. Here we included the reference genomes of *G. arboreum*, *G. hirsutum*, *B. vulgaris,* and *Arabidopsis thaliana* to define genes that were previously well-annotated. This analysis resulted in 28,022 orthogroups (OGs) with genes of one or more species and only 6,370 OGs were found to include genes from all five species. We define here orthogroups containing DEGs in response to pH treatment as DEG orthogroups. Gene expression of DEG orthogroups are shown in Supplementary Data, where we can see most of the DEG orthogroups showed a different expression pattern in each species. DEG orthogroups related to hormone response, calcium sensing, and photosynthesis were found to be differentially expressed in all species but differing in the specific isoforms.

### GO term enrichment analysis

To determine which key plant metabolic pathways are involved in sensing or responding to wetness and pH changes, we performed a gene ontology (GO) enrichment analysis with all the DEGs per species using the ClueGo package (Bindea et al., 2009) in Cytoscape (Otasek et al., 2019). We observed the GO term enriched by treatment comparison against the Dry control (Figure 3A, B and C) and Wet control (Figure 3D). We found no metabolic categories enriched in the DEGs in the pH4vsWet comparison. Response to hypoxia was a term shared across species and shared in all the three comparisons against the dry control (Figure 3A, 3B, 3C). We observed unique responses in every comparison shared across species (Figure 3, gray circles); furthermore, the pH4vsDry comparison (Figure 3B) is the stress condition that reveals the enrichment of the most signaling pathways in response to many different abiotic stresses such as salt stress, osmotic stress, oxidative stress, and to “external stimulus”, and to biotic stresses including “other organism” and fungus. In the pH2vsDry comparison (Figure 3C), cellular response to abiotic stimulus is driven by *G. hirsutum,* but we still observed response to osmotic stress and to light stimulus shared across species. In the pH2vsWet comparison (pH 6.5) we can separate the response of the leaf to pH changes, excluding the response to wetness (Figure 3D). Auxin transport and other metabolic processes involved in growth and development are impacted by pH stress together with photosynthesis-related pathways. On the other hand, we can observe *G. arboreum* (Figure 3) and *N. bicalcarata* (Figure 3) drove some of the GO term categories in WetvsDry (Figure 3A) and pH2vsWet (figure 3D) comparisons; while the pH4vsDry (Figure 3B) comparison is enriched mainly by DEGs from *N. bicalcarata* (Figure 3, yellow circles) and pH2vsDry (Figure 3C) comparison by *G. hirsutum* (Figure 3, pink circles).

**Figure 3.**
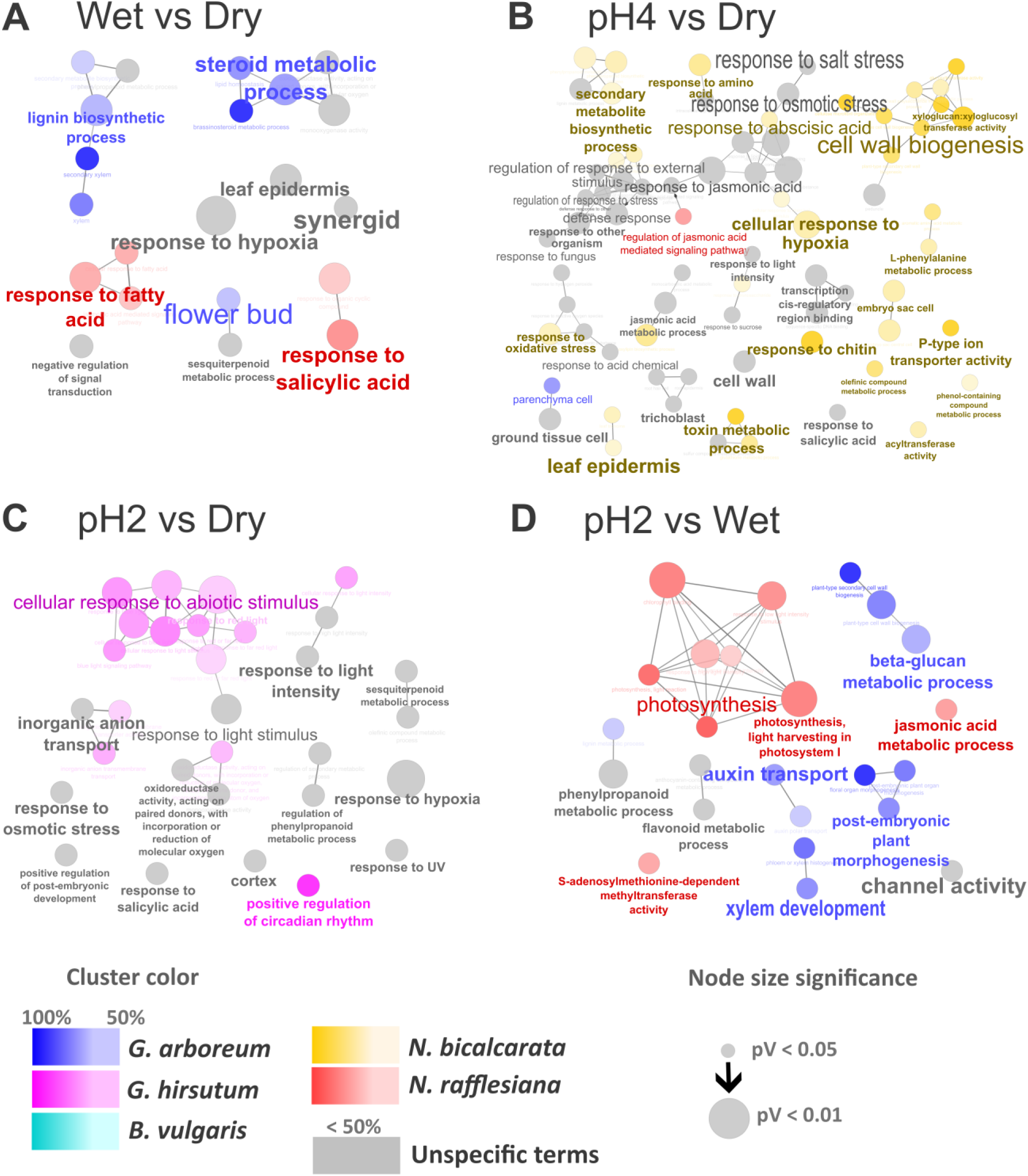
Responses to abiotic stresses are activated in response to wetness and pH. GO term enrichment of DEGs of the comparison of A) WetvsDry; B) pH4vsDry; C) pH2vsDry and D) pH2vsWet. Grey circles are GO terms shared across different species. Circles with different colors are GO terms that are enriched with more than 50% of the DEGs from an specific species found in the legend. The size of the circles represents how significant they are, the smallest started from a p-value of 0.05. “Response to hypoxia” was a GO term that was enriched in all species in response to all treatment comparisons. “Photosynthesis” GO term was enriched in pH2vsWet, especially for *N. rafflesiana*.

We visualized DEG orthogroups with Mapman (Thimm et al., 2004) to look for a gene expression pattern related to the pH gradient (Supplementary data, Figure 4). A general overview is shown in Suplementary Figure 1B, where we highlight in *G. hirsutum* how some genes in light reactions pathways are upregulated; however, following the pH gradient (from more neutral to more acidic) reveals that there are many others that were downregulated with decreasing pH. Also, in *N. bicalcarata,* photorespiration is the main metabolism where we observed an upregulation in the comparison against Dry control but not against Wet control (Supplementary Figure 1D).

We selected only the main metabolic categories that showed an interesting expression pattern from the metabolism overview, including external stimuli, phytohormones, solute transport and photosynthesis (Figure 4, Supplementary Data). As previously demonstrated, there are not many DEGs shared across species, but rather we observe that *G. hirsutum* and both *Nepenthes* species each have more DEGs in each metabolism category (Figure 4A, Supplementary Data). *G. hirsutum* visually shows increasing downregulation of DEGs moving down the pH gradient (Figure 4), mostly in solute transport and photosynthesis; this last one involves many genes related to the photosynthetic machinery such as the light harvesting complex (LHC) and the photosystem II assembly (PS-II). In contrast, *N. bicalcarata* has more upregulation of DEGS, which is mainly in the treatment comparison against the Dry control. In *N. rafflesiana,* more downregulation of DEGs related to photosynthesis and more upregulation of solute transport and phytohormone DEGs shows a pattern related to the pH gradient (Figure 4A).

**Figure 4.**
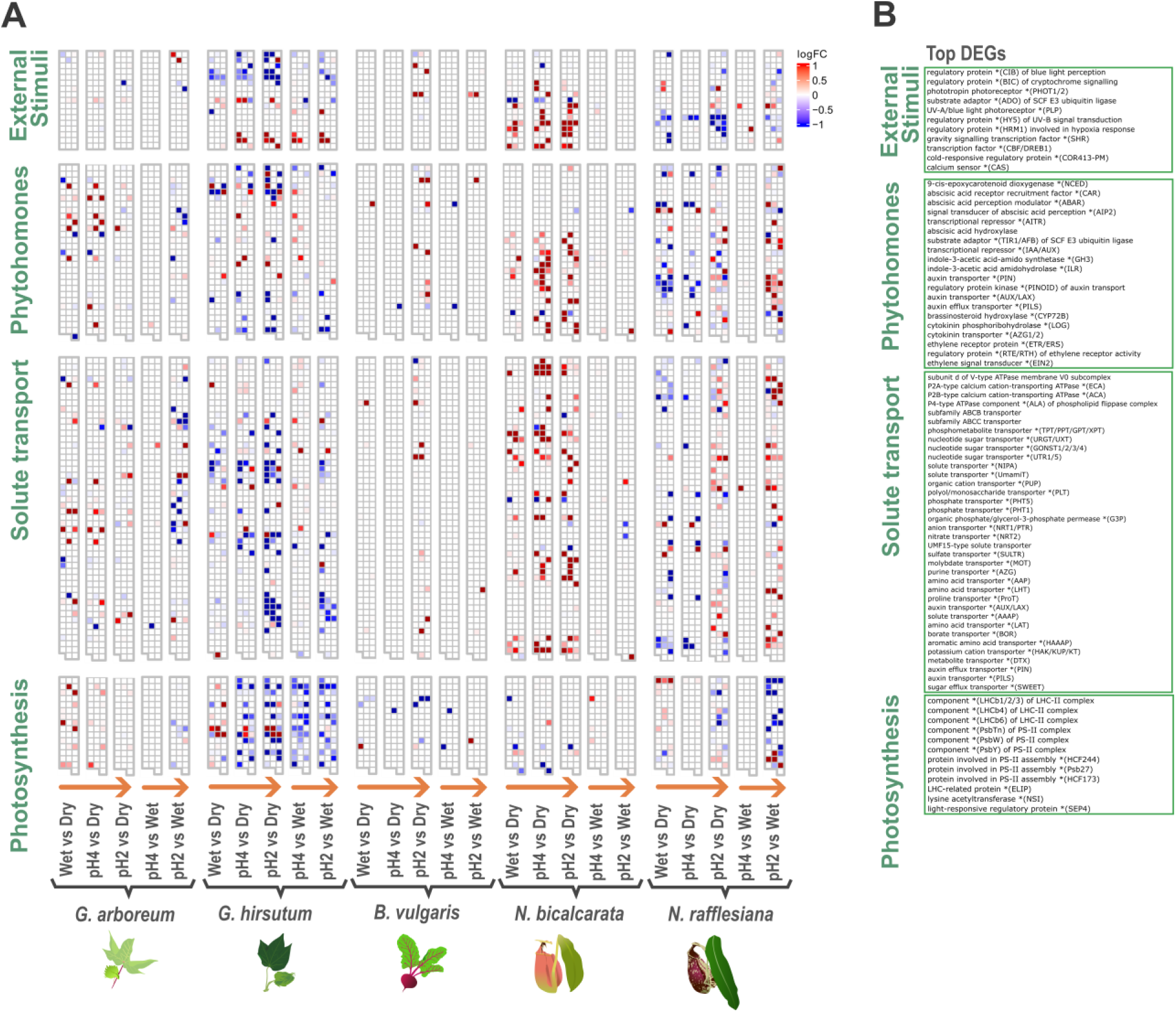
Photosynthesis is downregulated in response to pH stress. A) Expression pattern of orthologous DEGs across species and treatment comparison; orange arrows represent the increase in acidification response. B) Top 20 genes in each category. Orthogroup DEGs upregulated (red) and downregulated (blue). Notably, genes related to photosynthesis show a response to the pH gradient in *G. hirsutum*.

### Co-expressed orthogroups across species

To test if there was a cluster of co-expressed genes shared across species that can be involved in the response to pH changes, we used the orthogroups matrix from OrthoFinder and then we ran a cluster analysis by Clust (Abu-Jamous and Kelly, 2018). Only one cluster of 17 co-expressed orthogroups was found across species (Figure 5). A possible gene expression pattern driven by pH can be observed in both *Gossypium* species, but a mild effect is seen in both *Nepenthes* species, whereas no clear pattern is observed in *B. vulgaris* (Figure 5A). Genes in this cluster are shown in Figure 5B and we found genes related to the photosynthetic machinery (light harvesting complex, LHC) and calcium sensors (CAS). In *G. arboreum* we observed more upregulation of this cluster, which shows the opposite pattern of the one observed in *G. hirsutum* (Figure 5B). No clear pattern of fold change was observed in *B. vulgaris* or either *Nepenthes* species.

**Figure 5.**
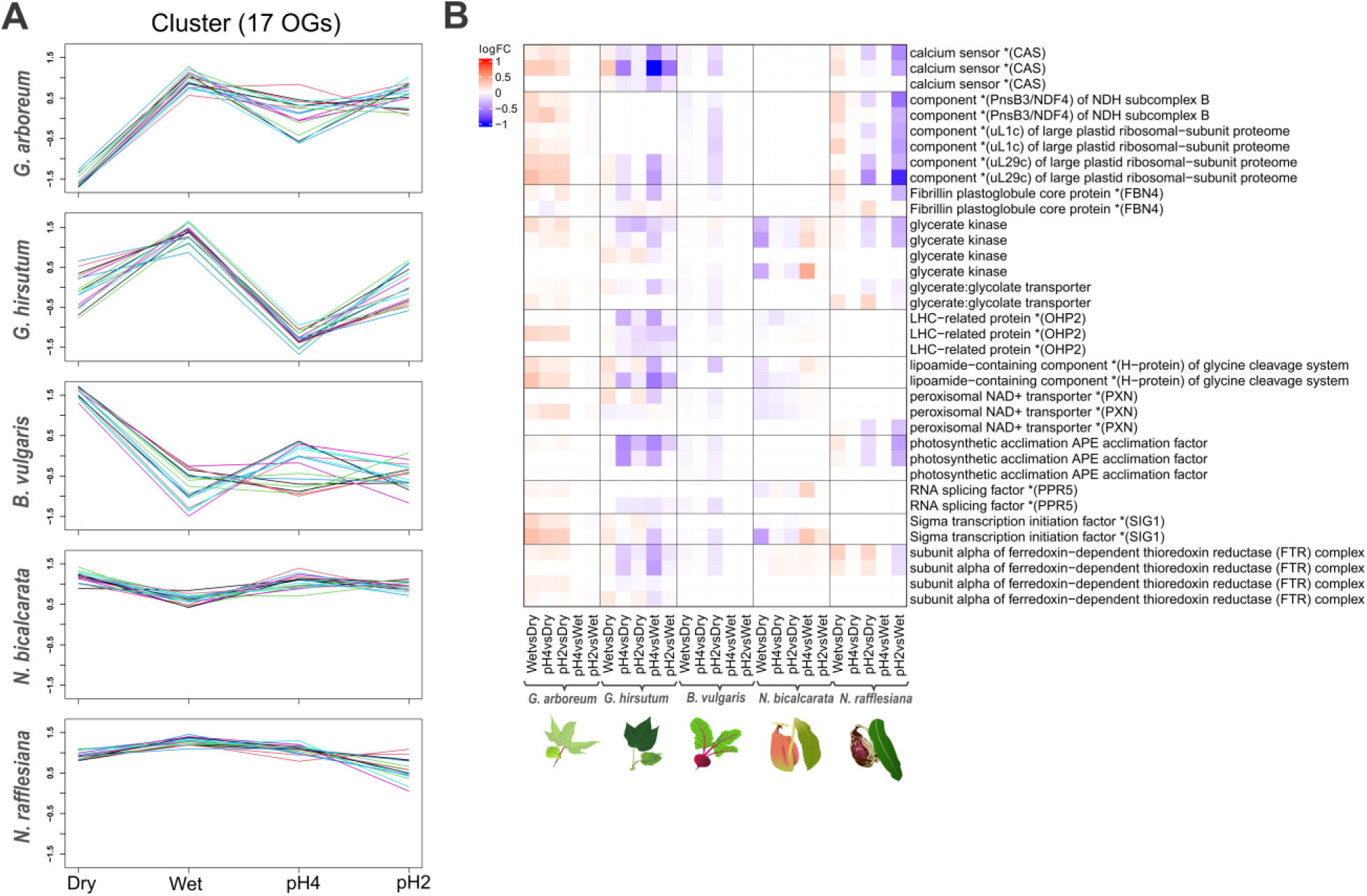
Light harvesting complex (LHC) and calcium sensor (CAS) genes are conserved across species and impacted by external pH changes, mostly in Gossypium species. A) Orthogroup expression patterns of a sinlge cluster that includes 17 orthogroups conserved across species in response to pH treatments. B) Expanded view of gene expression pattern of the cluster presented in A. The pattern of gene expression shows how they are differing by species.

## Discussion

### Phylloplane buffering ability is species-specific

The phylloplane is the main front line for sensing external stimuli and abiotic and biotic interactions for defense and to maintain homeostasis (Gilbert and Renner, 2021; Shepherd and Wagner, 2007). We were able to observe that external pH changes had a variable impact on the phylloplane of different plants (Figure 1). In the comparative transcriptomic analysis, all plants showed different numbers of DEGs in different pH treatment comparisons (Figure 2). A similar pattern of interspecific dissimilarity was found in a comparative transcriptomic analysis across *Arabidopsis thaliana*, *Hordeum vulgare* and *Oryza sativa* in response to different abiotic stresses including response to hormones such as abscisic acid and salicylic acid (Hartmann et al., 2022).

Not only can species differ in the composition of DEGs, but they can be linked to functional differences as well. In our study, many of these GO terms were driven by specific individual species (Figure 3); i.e., “secondary xylem” in lignin biosynthetic process and “brassinosteroids metabolic process” are enriched in *G. hirsutum* while “response to fatty acid” and “response to salicylic acid” were enriched in *N. bicalcarata* (Figure 3A). Additionally, a CLC chlorine sensor was found upregulated in *B. vulgaris* and *N. rafflesiana* in the pH2vsWet comparison, which suggests that these two species were able to detect the excess of Cl^-^ ions from the pH treatment. Altogether, even when both *Gossypium* species seem to buffer pH similarly (Figure 1A, B), and have similar number of total DEGs, their number of DEGs varies across treatment comparisons (Figure 2); this mismatch of the pattern of expression is also observed in both *Nepenthes* species, again showing how molecular mechanisms sensing and responding to external pH disturbances may vary with species identity.

These differences at the molecular level may translate to phenotypic differences across plants that impact the response to pH treatments at the organismal level. One likely relevant phenotypic consideration relates to differences in leaf morphological protective mechanisms (waxy cuticles, trichomes). Cuticles play an important role in transpiration, loss and uptake of polar solutes, and biotic interactions (Hauser’ et al., 1993; Zeisler-Diehl et al., 2018). Harr & Guggenheim (1995) described some properties of leaf surfaces in different crop species, including the contact angle of water on the leaf surface (e.g., contact angle = 34° for *G. hirsutum*, 65° for *B. vulgaris*) and the chemical composition of cuticular waxes (gas chromatography peaks); highlighting the difference of the chemistry of the cuticle between *B. vulgaris* and *G. hirsutum* if we compare their chromatography fingerprint, noticing 4 peaks with two major peaks at retention time around 5 and 7 in *B. vulgaris,* while in *G. hirsutum* there are 6 peaks but less abundant (Harr and Guggenheim, 1995). The composition of waxy cuticles can make leaf surfaces impermeable to water and dissolved solutes, and control how leaf surfaces can interact with microorganisms, even having a role in plant defense responses (Wang et al., 2020; Zeisler-Diehl et al., 2018). In our results, we found *B. vulgaris* with the lowest number of DEGs responding to pH stress (Figure 2). As these aforementioned leaf surface properties are fixed developmental differences, the different pH responses between *B.* vulgaris and *Gossypium* species are not necessarily only due to active response mechanisms. Finally, both *Nepenthes* species showed the highest number of DEGs, suggesting that these species possess a broad molecular mechanism to be aware of their surrounding environment (Figure 2, 4). Unlike the other species in this study, the *Nepenthes* were constantly subjected to a high humidity environment (80% RH), reflecting their native tropical habitat. More moisture can lead to an increase in biotic interactions with microbes and insects (Evans et al., 2022; Lebrija-Trejos et al., 2023), which might force the plants to develop more complex/reactive mechanisms for defense and sensing the surroundings. For instance, one of the top GO terms for *N. bicalcarata* was “response to other organism” and for *N. rafflesiana* “regulation of defense response” (Supplementary Figure S1). Further studies are needed to characterize deeper anatomical and chemical properties of leaves that may explain why, for some plants, it may be more relevant to buffer external pH changes, and for others, their physical/chemical leaf properties may be enough to cope with external disturbance.

### Ca^+^ role on pH stress response

The transport of a wide diversity of solutes is one of the basic metabolic processes involved in how plants sense their external surroundings. According to the metabolic categories identified by Mapman analysis, solute transport seems to be highly active, and differentially regulated across the pH gradient of increasing acidity, mainly in *G. hirsutum*, *N. bicalcarata* and *N. rafflesiana* (Figure 4). Inorganic elements (calcium Ca^2+^, potassium K^+^, phosphate P), organic compounds (sugar, amino acids) and hormones (auxin) transporters were found differentially expressed (Figure 4b). Ca^2+^ signaling pathway is the major point of signaling crosstalk between abiotic and biotic stresses (Dong et al., 2022; Li et al., 2022; Xu et al., 2022). An active metabolism involving Ca^2+^ ATPases and calcium sensors seems to be active in most of our plants (Figure 4b, Figure 5b). To our knowledge, the relationship of Ca^2+^ and pH homeostasis has not been widely studied. But studies of drought and salt stress have showed a relationship between pH and Ca^2+^; while levels of Ca^2+^ increase in the cytosol there is a concomitant decrease in pH (Feng et al., 2016; Gao et al., 2004; Ruan et al., 2024). More recently, it has been observed calcium transients are accompanied by pH changes in the cytosol in response to wounding, extracellular ATP or feeding a synthetic auxin (Behera et al., 2018).

In addition to the presence of intracellular Ca^2+^, these ions may also be transported to the external environment on the phylloplane. Salt-excreting glands or hydathodes in cuticles are known to modulate the loss or uptake of solutes (Zeisler-Diehl et al., 2018). In some species of *Gossypium*, glandular trichomes can store and excrete different ions such as calcium, magnesium and potassium onto the leaf surface (Elleman and Entwistle, 1982; Harr et al., 1984). Alkaline phylloplane pH is an extended phenotype found throughout the Malvales, which has not been observed in any other order (Gilbert and Renner, 2021). Some calcium-related genes were found to be differentially expressed in response to pH, such as the vacuolar cation/proton exchanger (CAX) which was downregulated in all treatment comparisons in *G. hirsutum,* and calcium sensors that were found in the conserved cluster across species. CAX antiporters are involved in the transport of Ca^2+^ or other cations using the H^+^ or Na^+^ gradient generated by primary transporters (Pittman and Hirschi, 2024; Shigaki et al., 2006). But more related to pH regulation is the presence of a V-type H^+^ ATPase pump which was upregulated in pH4vsWet comparison in *G. hirsutum* (Supplementary data); V-type proton ATPases are involved in pH regulation by primarily pumping H^+^ out of the vacuolar membranes (Liang et al., 2015; Müller et al., 1996). This also highlights the role of the vacuoles in the regulation of pH. Vacuoles are known to store different types of acid compounds to maintain the homeostasis of intracellular pH as well as playing a critical role in signaling pathways to abiotic or biotic stress (Faraco et al., 2014; Martinoia et al., 2018; Zhang et al., 2014). Additionally, we found a potassium transporter downregulated in the pH2vsWet comparison in *G. arboreum* (HAK2) and in the pH4vsDry comparison in *N. rafflesiana* (HAK5) while in *N. bicalcarata*, the transporter (HAK19) was upregulated in pH4 and pH2vsDry comparisons (Supplementary Data). The role of this K^+^ transporter in *Gossypium* could be related to an intracellular metabolism or the storage of K^+^ in the glandular trichomes.

### Photosynthesis is negatively impacted by pH stress

Photosynthesis is probably the main metabolic pathways that comes to mind when considering leaves. Genes involved in the photosynthetic machinery and response to light seem to be differentially expressed in response to pH, and are mostly downregulated following the increase in acidity, this is most apparent in *G. hirsutum* and *N. rafflesiana* (Figure 4). This is one of the few metabolic functions that are shared across species, and it is clearly negatively impacted by pH stress. Stroma, the site where the photosynthetic machinery is located, is known to be susceptible to changes in intracellular pH (Aranda Sicilia et al., 2021). Acidification of the stroma generally leads to a disruption of the photosynthetic machinery involving the photosystem and the electron chain, which will be followed by a chlorophyll breakage (Tikhonov, 2012; Trinh and Masuda, 2022). Light harvesting complex and photosystem II genes were mainly downregulated in all of our species (Figure 4, Supplementary Data). This is probably to avoid cell damage due to acidification.

The negative impact of pH stress to photosynthesis can have repercussions from an ecological point of view on how the landscape will be shaped due to increase in external acidification agents such as acid rain, foliar pesticides and fertilizers. Thus, it may be beneficial to study how external acidification can impact crops of commercial interest.

### The pH molecular response is interlinked to other abiotic and biotic stresses signaling pathways

In nature, plants are more likely to interact with a combination of stressors, thus plant response to multiple stresses can be interconnected in a major signaling pathway to acclimate fast and more efficiently in a changing environment (Zandalinas et al., 2020). Response to different types of abiotic and biotic stresses have been seen to converge at some point in their molecular response pathway (Li et al., 2022; Munns and Millar, 2023; Pereira, 2016; Zandalinas et al., 2020; Zhang et al., 2023). Here we found DEGs related to different abiotic and biotic responses mostly activated in response to the pH4 treatment and driven mainly by *Nepenthes* (Figure 3). This suggests that leaves sensing external mild pH activates other abiotic stresses or the general response to abiotic stress in which pH is integrated as a stressor. One of the few shared GO terms enriched across species was related to “hypoxia” in all the comparisons against Dry control (Figure 3A, 3B, 3C), suggesting that all species were sensing wetness (i.e., slight reduction in oxygen on the surface when covered in water droplets). Sensing wetness on the leaf surface seems to activate biotic responses as we can infer from the GO term “interaction with other organism” and “response to fungus” (Figure 3b). Plant response to biotic stress triggers the production of salicylic acid, jasmonic acid and ethylene depending on the pathogen (Cohen and Leach, 2019), which were also GO terms enriched in our results (Figure 3). Understanding the ecological implications of this buffering ability in the interactions with other organisms such as microbes and insects seems to be highly relevant.

Furthermore, plant responses to abiotic stress such as drought, cold, salinity and wounding share a similar signaling pathway backbone (e.g. genes, molecules) (Kidokoro et al., 2017; Skalak et al., 2021; Verma et al., 2016). Perception of these stresses starts with membrane receptors (GPCRs, G protein-coupled receptors; RKLs, receptor-like kinases; histidine kinases; and ion channels). These receptors lead to changes in the pool of Ca^2+^ which activates several kinases, such as CDPKs (calcium-dependent protein kinases). Nevertheless, even when the plant uses similar players for different stresses, the final section of the response mechanism is more specific to the type of stress.

## Conclusions

Comparative transcriptomic analyses of leaf responses to external pH changes showed how pH stress impact differs between species. Mild pH changes trigger the activation of signaling pathway genes involved in the response to many other abiotic stresses such as osmotic stress, drought stress, etc. Furthermore, we observed pathways related to interaction with other organisms, and defense was turned on due to wetness and mild pH. *Gossypium*, as the genus able to alkalinize acidic pH, involves the participation of Ca^2+^ and H^+^ ATPases. Further studies are needed to characterize these enzymes in other species in *Gossypium* to clarify the underlying molecular mechanism and determine how this could be interlinked to the general abiotic stress backbone.

## Materials & Methods

### Biological samples

We acquired seeds of *G. arboreum* (accession PI 615701), *G. hirsutum* (accession PI 529181), and *B. vulgaris* (accession Ames 3060) from the United States Department of Agriculture (USDA) Germplasm Resources Information Network (GRIN). The seeds were planted in Sunshine #4 Soil Mix (insert brand/supplier) and grown in the Pennsylvania State University Department of Entomology headhouse greenhouse under ambient conditions. Because it was not feasible to grow *Nepenthes* from seed, we attained small adult plants (clonally propagated) of *N. bicalcarata* (accession BE-3031) and *N. rafflesiana* (accession BE-3722) from Borneo Exotics ltd. Supplied by Carnivero (Austin, TX, USA). The *Nepenthes* were potted in a sphagnum medium and grown in a Conviron reach-in growth chamber with the following settings: 12-hour day-night cycle, daytime temp = 28°C, nighttime temp = 26°, relative humidity = 80%, fluorescent/ incandescent levels = 250 uMol. The *Nepenthes* pots were also treated with imidacloprid to clear a scale infestation before the start of the experiment. For the plants grown from seed, they were all planted in October 2020 and grown until all species had germinated, emerged, and produced the first two nodes of true leaves (4-6 weeks from being sown). We aimed to perform all the pH manipulation experiments on the same day for these youngest fully expanded leaves to minimize temporal/ontogenetic variation; this greenhouse experiment and subsequent harvest occurred on 14 November 2020. The *Nepenthes* lamina experiment and harvest in the growth chamber occurred on 10 December 2020.

For the pH manipulation experiments, the leaf was sprayed with one of three pH treatments: pH 6.5, 4.0, and 2.0. These sprays were created by mixing distilled deionized water (ddH2O) with the appropriate volume of HCl to reach the target pH level. These pH solutions were stored in inert glass spray bottles to avoid leaching of material or change in pH due to CO2 diffusion. The leaf was sprayed until thoroughly wet (∼15 sprays) and collected directly into 2.0 mL DNAse/RNAse-free tubes of RNALater after 5 minutes of exposure. To collect the leaves, we used autoclaved forceps wiped with RNAse Away, pulling the leaf from the abcission zone of the petiole, or in the case of *Nepenthes* lamina, the weak point of the stem *(Nepenthes* lamina experiments and harvesting took place after pitchers were removed days prior). The pH response of the leaf was monitored throughout the 5-minute exposure period using a flat-tipped pH probe (Hanna Instruments), and the final pH value was recorded. For our unmanipulated controls, we sampled dry (unsprayed) leaves into RNALater, and these dry pH levels were also measured and recorded. For each species, we sampled three replicates per treatment (including the dry control), where a separate individual potted plant is one replicate. The RNALater-stabilized samples were stored at 4 C until extraction.

### RNA extraction and Sequencing

We conducted total RNA extractions using the Spectrum Plant Total RNA kit, according to manufacturer’s protocol with some modifications. First, the leaves were transferred into clean tubes and centrifuged at max speed for 30 seconds to remove remnant RNALater. For the mechanical lysis step, we used a single large steel bead (autoclaved and sterilized with RNAse Away) to homogenize the leaf sample in lysis buffer using a FastPrep machine set to 5.5 m/s for 20 s. Following homogenization, the sample was incubated in a heatblock at 56 C for 5 minutes, and the remaining steps proceeded as per the original protocol. Our extracted RNA was sent to Novogene for library prep, quality control, and sequencing on an NovaSeq 6000 PE150 platform.

### Transcriptomic analyses

Each RNAseq analysis was performed for each species separately. The raw read quality of each paired-end library was tested by FastQC v0.11.9 (Andrews, 2010). Adapters and low-quality bases were removed with Trimmomatic (Bolger et al., 2014). Transcriptome *de novo* assembly was performed using Trinity v2.9.1 (Grabherr et al., 2011) with default settings for each species to treat them all the same. Reads counts were done with Salmon v1.5.0. Low counts (< 3 TPM) were filtered out and the most highly expressed isoform for each gene was kept. Transdecoder (Haas and Papanicolaou, 2017) was used to identify ORF coding region in all the assemblies.

Annotation was done with Trinotate v4.0.2 (Haas et al., 2013) and EnTAP v1.0.1 using Swissprot and PFAM protein database mainly. Salmon counts were filtered by taxonomy, only genes that were annotated for Viridiplantae taxa were considered.

### Differential gene expression analysis

In order to identify relevant genes involved in pH response differential gene expression (DGE) analysis was performed with DESeq2 v1.38.3 package (Love et al., 2017; Zhang et al., 2018) for each species separately. Batch correction for each species was performed with COMBAT_Seq v0.0.4 package (Zhang et al., 2018). DGE analysis were performed as follow: the sample taken without spraying any liquid was considered the dry control (Dry) and the pH 6.5 sample was considered the wet control (Wet) due to its close pH to the common distilled water pH.

DGE analysis was made comparing each pH treatment (Wet, pH 4 and pH 2) against the dry control (WetvsDry, pH4vsDry, pH2vsDry) and comparing pH 4 and pH 2 treatment against wet control (pH4vsWet, pH2vsWet). Differentially expressed (DE) genes were extracted with a cutoff of *logFC* > |1| and a *p-adjusted* value < 0.05 for each species. Graphic representation of the number of DE genes was done using ComplexUpset package v1.3.3 in RStudio v4.2.3. ClueGo v2.5.9 (Bindea et al., 2009) throught Cytoscape v3.10.2 (Otasek et al., 2019) was used for enrichment analysis of GO term.

### Orthologous groups across species and cluster analysis

Protein alignment from transdecoder protein pep file output for each species were used altogether with the protein sequences from the three reference genomes available (*G. arboreum, G. hirsutum,* and *B. vulgaris*) to find the orthologous match across species with OrthoFinder v2.5.4 (Emms and Kelly, 2019). The protein sequences of the references genome were downloaded from ENSEMBL for *B. vulgaris v1.2.2* (https://plants.ensembl.org/; accessed date: 02-20-2023) and from NCBI via FTP (https://ftp.ncbi.nlm.nih.gov/genomes/all/; accessed date: 02-20-2023) for *G. hirsutum* v2.1 (accession GCF_007990345.1), and *G. arboreum* v2 (accession GCF_025698485.1). The orthogroups ids file altogether with the gene expression dataset were used to perform a co-expression clustering analysis with Clust v1.18.0 (Abu-Jamous and Kelly, 2018) to identify co-expressed gene clusters across species.

### Statistical analysis

Statistical analysis such as one-way ANOVA, Dunnet test and LDS-Fisher and generalized linear model (glm) test were performed in RStudio v4.2.3.

## Acknowledgements

We acknowledge Scott Diloreto and Adam Bettinger for help with rearing plants in the greenhouse and growth chamber, as well as help from John Fulginiti and Elise Elizondo in preliminary methods testing. We also thank Chloe Drummond, Adam Rork, Kylie Bocklund, members of the Plant Resilience Institute, and the Gilbert Lab for helpful discussions. KJG was supported by USDA-NIFA grant #s 2019-67012-29872 and 2019-67012-37587. TR acknowledges support from NSF grant DEB 2030871. Any opinions, findings, conclusions, and recommendations expressed in this material are those of the authors and do not necessarily reflect the views of the National Science Foundation.

This work is also supported by the USDA National Institute of Food and Agriculture and Hatch Appropriations under Project #PEN04974 and Accession #7006543.

## Author contributions

KJG and TR initially conceived of the idea, KJG performed greenhouse/growth chamber experiments and conducted RNA extractions, CLG and KJG performed bioinformatics and data analyses. CLG wrote the initial draft and CLG, JBF, TR, and KJG contributed to writing the final version of the manuscript.

