## Supplementary figures and images for "Species-specific phyllosphere responses to external pH change"

### Supplemental Figure

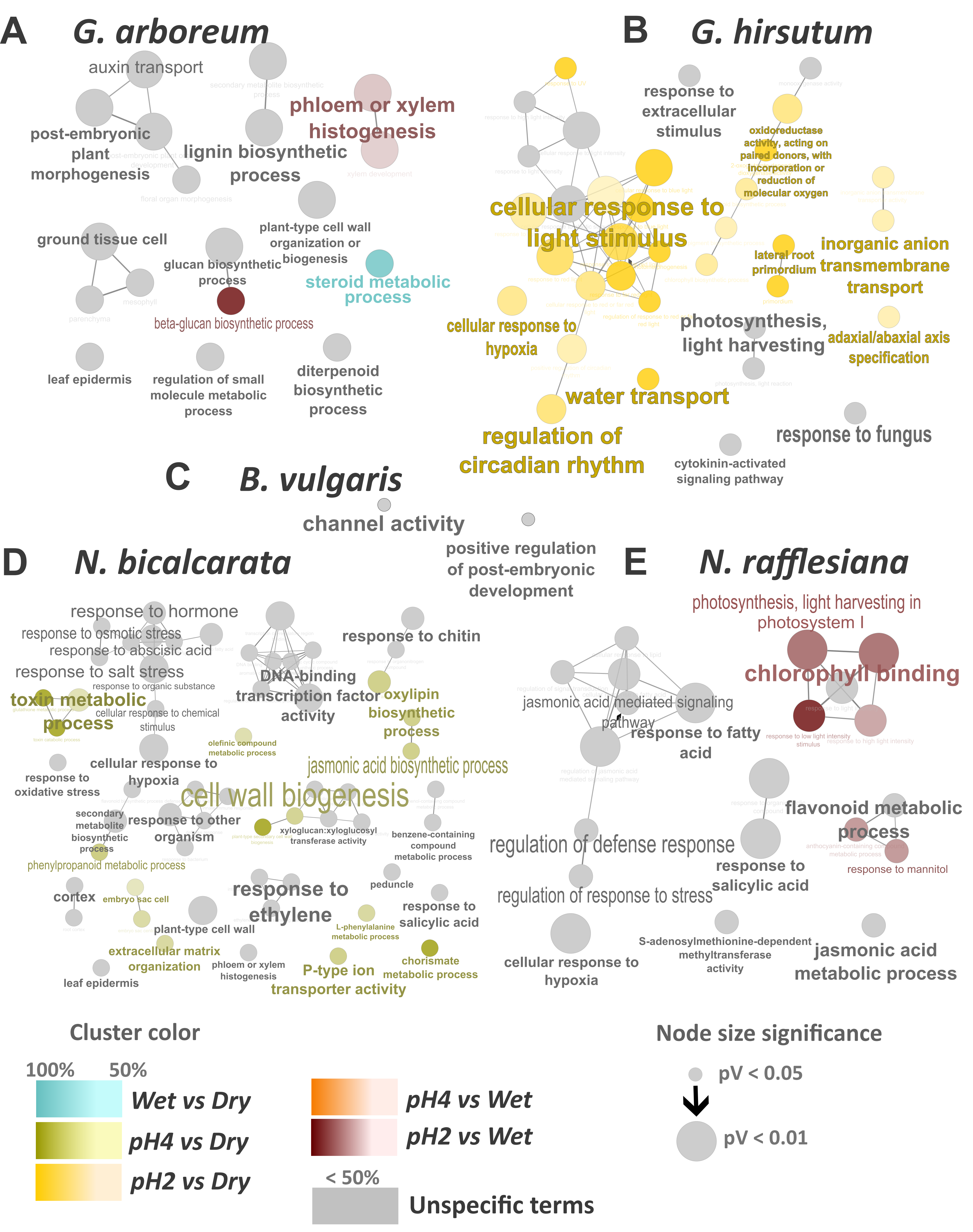
